# Astrocytic D1 Dopamine-Signaling Regulates Synaptic Remodeling and Cocaine Seeking

**DOI:** 10.64898/2026.05.13.724947

**Authors:** Shi Yan, Qize Li, Yao Wang, Wanqiu Chen, Zhong Chen, Alexander K. Zinsmaier, Paloma Huguet, Zhengsong Lu, Xiguang Qi, Zheng Xu, Yi Han, Gouri Sharma, Charles Wang, Eric J. Nestler, Oliver M. Schlüter, Yan Dong

## Abstract

D1-type receptor (D1R)-mediated dopaminergic signaling within the nucleus accumbens shell (NAcSh) is essential for forming adaptive circuit changes that embed persistent memories associated with drug seeking and craving. While D1R is expressed in both NAcSh neurons and astrocytes, the specific contribution of astrocytic D1R to drug-related circuit plasticity remains poorly understood. Here, we demonstrate in mouse NAcSh slices that D1R agonists increase astrocytic Ca^2+^ activity, an effect that is attenuated by astrocyte-specific D1R knockdown. Furthermore, selective knockdown of astrocytic D1R in the NAcSh prior to cocaine self-administration inhibits cocaine-induced generation of silent synapses, therefore hampering the associated remodeling of NAcSh circuits. Behaviorally, knockdown of NAcSh astrocytic D1R expedites the extinction of cocaine seeking and reduces cue-induced reinstatement. These results identify astrocytic D1R as a fundamental component of NAcSh dopamine signaling during cocaine experience that remodels synaptic connections and neural networks underlying drug seeking and relapse.

## Introduction

The nucleus accumbens shell (NAcSh) is a key brain region where drug-induced adaptations at glutamatergic synapses remodel neural circuits, ultimately driving drug craving and seeking associated with substance use disorder (SUD) (1, 2). A hallmark of this remodeling is the generation of AMPA receptor (AMPAR)-silent glutamatergic synapses (3, 4). While silent synapses are primarily enriched in the developing brain as nascent synaptic contacts that initiate new neural networks, their levels decline substantially after circuit maturation (5–10). In adult animals, however, cocaine self-administration generates new silent synapses on NAcSh medium spiny principal neurons (MSNs) (11, 12). During drug abstinence, a subset of these synapses matures by recruiting AMPARs, thereby stabilizing new synaptic contacts in the NAcSh (11, 13, 14). Through the de novo generation and subsequent maturation of silent synapses, cocaine experience remodels the connectivity pattern of NAcSh circuits to support the behavioral responses associated with drug craving and relapse (2, 15). While mounting evidence suggests that cocaine-induced generation of silent synapses in the adult NAcSh requires the reactivation of developmental mechanisms involving astrocytic signaling (8, 12, 16, 17), how cocaine experience initiates this astrocyte-mediated synaptogenesis remains poorly understood.

Cocaine self-administration robustly elevates extracellular levels of dopamine in the NAcSh. Dopamine actions are predominantly mediated by its receptors, among which the D1 type receptor (D1R) is essential for several cocaine-induced molecular, cellular and behavioral adaptations. In the NAcSh, D1R is primarily expressed in two major cell types: MSNs and astrocytes. Disrupting NAc D1R-signaling impairs both cocaine-induced molecular and cellular adaptations in the NAcSh and SUD-associated behaviors (18–21). However, because most of these findings are obtained through nonspecific manipulations of D1R that do not distinguish between neuronal versus non-neuronal locations, the specific role of astrocytic D1R in cocaine-induced circuit and behavioral alterations remains unchecked.

Focusing on astrocytic D1R in mice, our current study shows that D1R agonists increase astrocytic Ca^2+^ activity in NAcSh slices, an effect attenuated upon astrocyte-specific D1R knockdown. Furthermore, knocking down D1R in NAcSh astrocytes prior to cocaine self-administration dampens cocaine-induced generation of silent synapses. Behaviorally, while disrupting astrocytic D1R-signaling does not affect the acquisition of cocaine self-administration, it expedites the extinction of operant responses for cocaine and reduces cue-induced reinstatement of cocaine seeking. These results establish a critical role of astrocytic D1R-signaling in silent synapse-mediated remodeling of NAcSh circuits and SUD-associated behaviors.

## Results

### Astrocytic D1R is targeted by cocaine

Evidence from immunostaining, mRNA hybridization, or cellular responses to receptor-selective ligands suggests the existence of both D1 and D2 dopamine receptors on astrocytes in the NAcSh and dorsal striatum (22–26). To determine the transcriptional impact of cocaine on astrocytic D1R, D2R, and other signaling components, we performed single-nucleus RNA sequencing (snRNAseq) of NAcSh cells from mice after 20 days of abstinence from cocaine or saline (control) self-administration.

To reduce inter-individual variability, we pooled NAcSh tissue from six mice per experimental group and performed droplet-based snRNA-seq (10x Genomics). Collectively, 11,029 nuclei from the saline group, and 17,611 nuclei from the cocaine group were captured and passed quality control, with an average of 2,280 detected genes per nucleus. Uniform manifold approximation and projection (UMAP) and unsupervised clustering analyses revealed 17 clusters based on their gene expression profiles, with cell types annotated using established marker genes. Astrocytes were readily distinguishable from D1R- and D2R-expressing MSNs and other cell types, showing minimal systematic shift in global cell identity between the cocaine and saline groups (Fig. 1A, B). In the astrocyte cluster, we detected Drd1 and Drd2 mRNAs. Furthermore, the cocaine group exhibited a higher average Drd1 expression level and a greater proportion of Drd1+ astrocytes, suggesting an upregulation of Drd1 transcription in NAcSh astrocytes following cocaine self-administration and abstinence (Fig. 1C).

**Figure 1.**
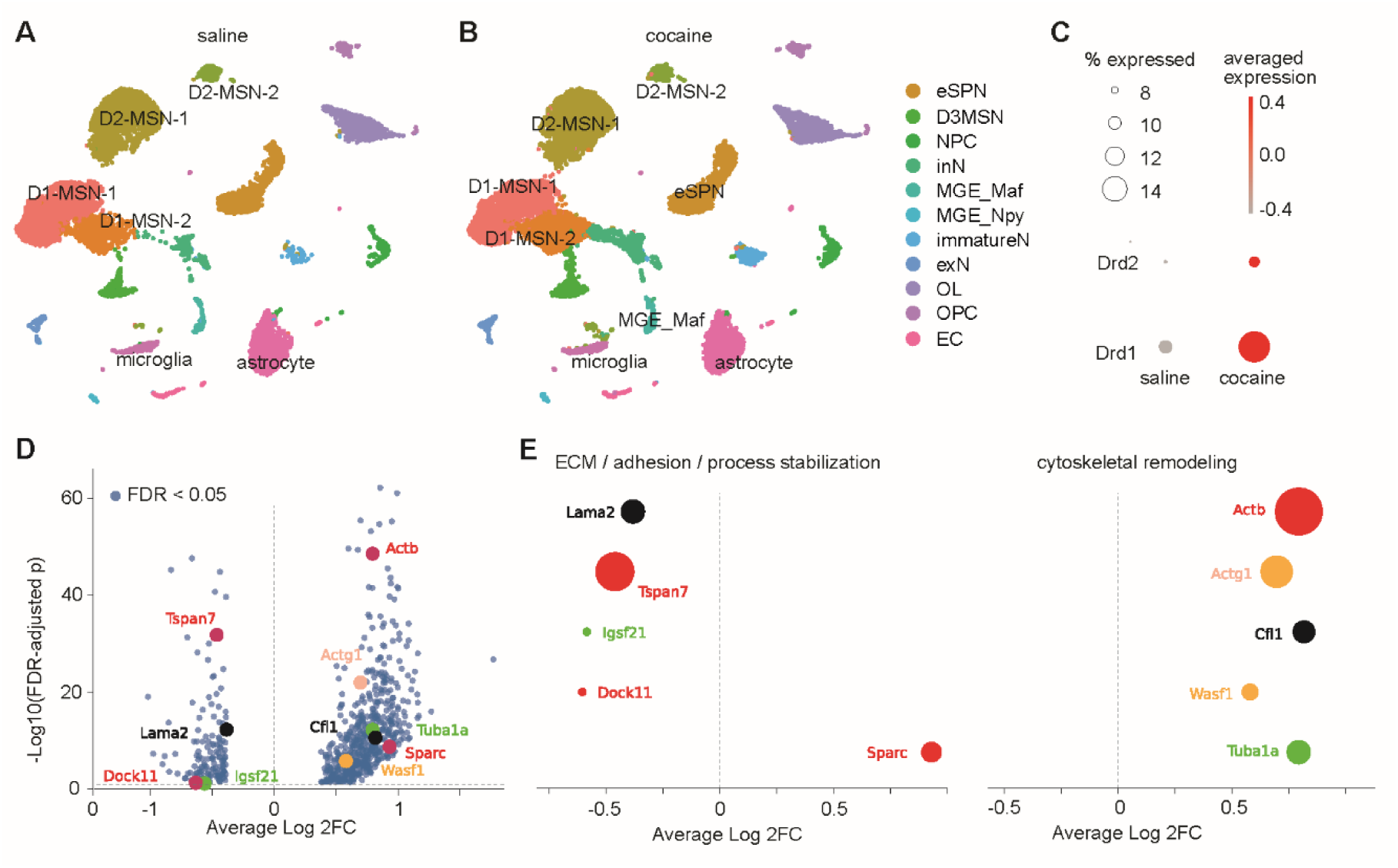
Transcriptional upregulation of astrocytic Drd1 following cocaine abstinence. (A, B) UMAP visualization of single nuclei from the NAcSh of mice following saline (A) or cocaine (B) self-administration and abstinence. (C) Dot plot showing increased proportion of Drd1+ astrocytes and increased average Drd1 expression in NAcSh astrocytes following cocaine self-administration and abstinence. (D) Volcano plot showing differentially expressed genes (log2FC > 0.38, FDR < 0.05) in astrocytes following cocaine self-administration compared to saline self-administration. X-axis indicating average Log2 fold changes (Average Log2FC), with positive vs. negative values representing upregulation vs. downregulation in cocaine mice relative to saline mice. Y-axis indicating −Log10 (FDR-adjusted *p* values). Genes outside the display range not shown. Ten morphology-related genes of interest are highlighted and labeled in corresponding colors. Dashed lines indicating Average Log2FC = 0 and FDR-adjusted *p* = 0.05. (E) Select astrocytic DEGs associated with reduced territory or process complexity following cocaine self-administration. Genes grouped by functional relevance to ECM / adhesion / process stabilization (left) or cytoskeletal remodeling (right). Dot sizes indicating the values of −Log10 (FDR-adjusted *p* values), X-axis indicating Average Log2FC of cocaine mice relative to saline mice.

Next, we performed differential gene expression analysis of astrocyte nuclei between cocaine and saline groups. Collectively, 558 genes were upregulated and 149 genes were downregulated in mice that self-administered cocaine (log2FC > 0.38, false-discovery rate (FDR) < 0.05) (Fig. 1D). Among these differentially expressed genes (DEGs), several were relevant to astrocyte morphology and process organization. Specifically, extracellular matrix (ECM)-related genes Lama2, Tspan7, Igsf21, and Dock11 were downregulated, while Sparc was upregulated (Fig. 1D, E), suggesting an altered structural support and process stabilization in cocaine mice. In parallel, cytoskeletal remodeling-related genes Actb, Actg1, Cfl1, and Tuba1a were upregulated (Fig. 1D, E), suggesting increased dynamics in astrocyte morphology. These DEG profiles predict dynamic morphometric changes in NAc astrocytes following cocaine self-administration (see below).

### Astrocyte-specific knockdown of D1Rs

To selectively manipulate D1R in astrocytes, we used an adeno-associated virus vector (AAV) with the AAV5 serotype, driven by an astrocyte-specific promoter (27, 28), a strategy previously validated for targeting NAcSh astrocytes (12). We designed an shRNA targeting the Drd1 transcript with a knockdown efficacy of ∼60% (Fig. S1). We co-expressed it with a green fluorescence protein (GFP) reporter. Four weeks following intra-NAcSh injection of the AAV2/5-GFP or AAV2/5-GFP-shD1R, we observed extensive GFP expression (Fig. 2A). These GFP+ cells exhibited characteristic ‘bushy’ morphologies, consistent with astrocytes (Fig. 2A, B).

**Figure 2.**
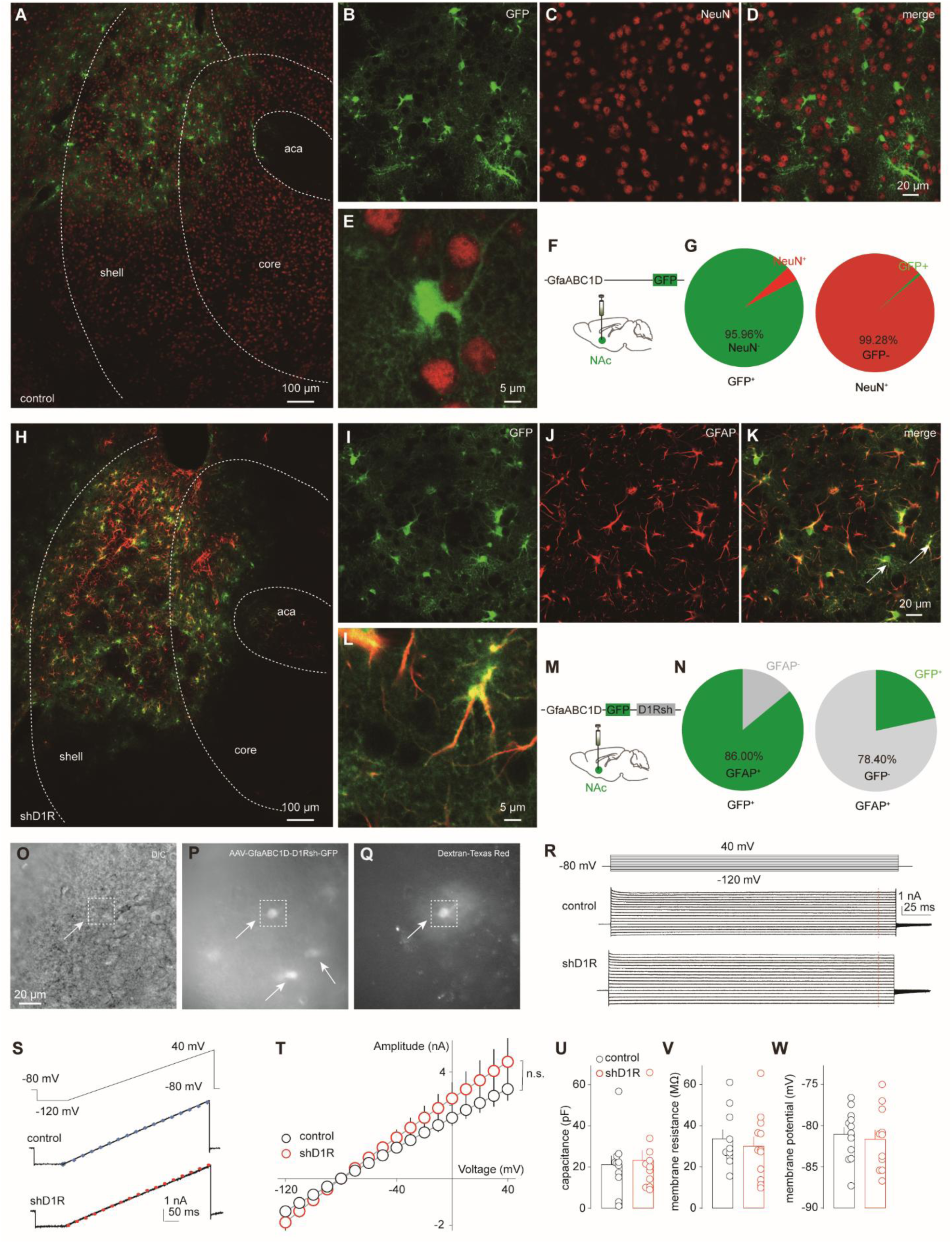
Knockdown of D1R does not affect the electrophysiological properties of NAcSh astrocytes. (A) Confocal image showing GFP expression following injection of AAV5-GfaABC1D-GFP into the NAcSh. (B-D) Representative images showing GFP-expressing cells (B), NeuN immunoreactivity (C), and merged images (D) in the NAcSh. (E) High-magnification image illustrating an example GFP-positive cell lacking NeuN immunoreactivity. (F) Schematic of virus injection for experiments shown in (A-G). (G) Summaries showing the proportion of GFP-positive cells that were NeuN-positive (left) and the proportion of NeuN-positive cells that expressed GFP (right). (H) Confocal image showing GFP expression following intra-NAcSh injection of AAV5-GfaABC1D-GFP-shD1R. (I-K) Representative images showing GFP (I), GFAP immunoreactivity (J), and merged signals (K), demonstrating colocalization of GFP with GFAP-positive astrocytes (indicated by arrows). (L) High-magnification image illustrating GFP-positive cells co-labeled with GFAP. (M) Schematic illustrating viral injection for experiments shown in (H-N). (N) Summaries showing the proportion of GFP-positive cells exhibiting GFAP immunoreactivity (left) and the proportion of GFAP-positive cells expressing GFP (right). (O) Example differential interference contrast (DIC) image of NAcSh slice showing a recorded astrocyte (arrow; dashed box). (P) Fluorescence image of the same field of view showing GFP expression from AAV5-GfaABC1D-shD1R-GFP–infected cells, with the recorded cell indicated (arrow; dashed box). (Q) Fluorescence image showing intracellular Dextran–Texas Red loaded through the patch-clamp pipette, confirming precise recording of the GFP-positive cell. (R) Representative whole-cell currents from NAcSh cells expressing GFP (control) or shD1R-GFP during a voltage-step protocol (top). Cells were held at −80 mV and stepped from −120 to +40 mV. (S) Representative whole-cell currents recorded from GFP-versus shD1R-GFP-expressing NAcSh cells in response to a voltage-ramp protocol (top), in which the membrane potential was linearly ramped from −120 to +40 mV. Dots indicate current amplitudes measured at corresponding membrane potentials obtained from the voltage-step protocol shown in (R). (T) Summary of current–voltage (I–V) relationships for GFP-and shD1R-GFP-expressing NAcSh cells. Current amplitudes were quantified at each holding potential from the voltage-step protocol shown in (R) and averaged across cells. Data represent mean ± SEM (control, 1237± 383.6, n = 11/6; shD1R, 1615± 503, n= 12/6, main effect of group *F*_1,21_ = 1.0, p = 0.32, time × group interaction *F*_1,21.1_ = 1.1, p = 0.31, Two-way repeated-measures ANOVA Greenhouse–Geisser corrected). (U-W) Passive membrane properties of NAcSh cells expressing GFP versus shD1R-GFP, including whole-cell capacitance (in pF: control, 21.17 ± 4.4, n = 11/6; shD1R, 23.2 ± 5.0, n = 11/5, *t*_19.7_ = 0.3, *p* = 0.76, unpaired Welch’s *t* test; U), membrane resistance (in MΩ: control, 33.6 ± 4.5, n = 10/6; shD1R, 30.1 ± 4.6, n = 12/5, *t*_19.9_ = 0.5, *p* = 0.59, unpaired Welch’s *t* test; V) and resting membrane potential (in mV: control, -81.1 ± 0.9, n = 12/6; shD1R, -81.7 ± 1.2, n = 11/5, *t*_19.3_ = 0.4, *p* = 0.68, unpaired Welch’s *t* test; W). Each circle represents one cell; bars indicate mean ± SEM; *, *p* < 0.05; **, *p* < 0.01.

Additional immunohistochemical staining of NeuN, a neuronal marker, showed minimal overlap between GFP+ and NeuN+ cells, indicating that the shD1R expression was restricted from neurons (Fig. 2B-G). Conversely, dual staining revealed a high degree of colocalization between glia fibrillary acidic protein (GFAP)+ and GFP+ cells, suggesting the preferential expression of shD1R in astrocytes (Fig. 2H-N).

To determine whether D1R knockdown altered the fundamental physiological properties of NAcSh astrocytes, we performed whole-cell voltage-clamp recordings to compare the current-voltage (I-V) relationship between control-GFP+ and shD1R-GFP+ cells. In response to a series of voltage steps, both control-GFP+ and shD1R-GFP+ cells exhibited large, passive leakage currents, a hallmark electrophysiological feature of mature astrocytes (Fig. 2R). Furthermore, voltage ramp protocols elicited gradually increasing currents without the presence of fast-activating or inactivating conductances (Fig. 2S), further confirming their astrocytic identity. Steady-state currents, measured 10 ms prior to the end of 250-ms steps, were similar between control-GFP+ and shD1R-GFP+ cells, resulting in overlapping I-V curves (Fig. 2T). Collectively, these results demonstrate that our virus approach specifically targeted NAcSh astrocytes and that the basal membrane physiology remained unaffected by D1R knockdown.

### D1R knockdown dampens D1R-induced astrocytic Ca^2+^ activity

Unlike neurons, whose activity relies on electrical signals, astrocytes primarily respond to D1R activation and other external stimuli through intracellular Ca^2+^ dynamics. To assess these dynamics, we expressed the genetically encoded Ca^2+^ indicator GCaMP6f selectively in astrocytes and recorded Ca^2+^-mediated fluorescence signals in brain slices. Slices were perfused with either a bath control or the D1R agonist SKF81297 (1 µM), in the presence of tetrodotoxin (TTX, 1 µM) to eliminate action potential-dependent neuronal influence. Using the NoRMCorre algorithm (see Methods), we defined Areas of Interest (AOIs) as regions exhibiting fluorescence transients during the 3-min recording period (Fig. 3A-C).

**Figure 3.**
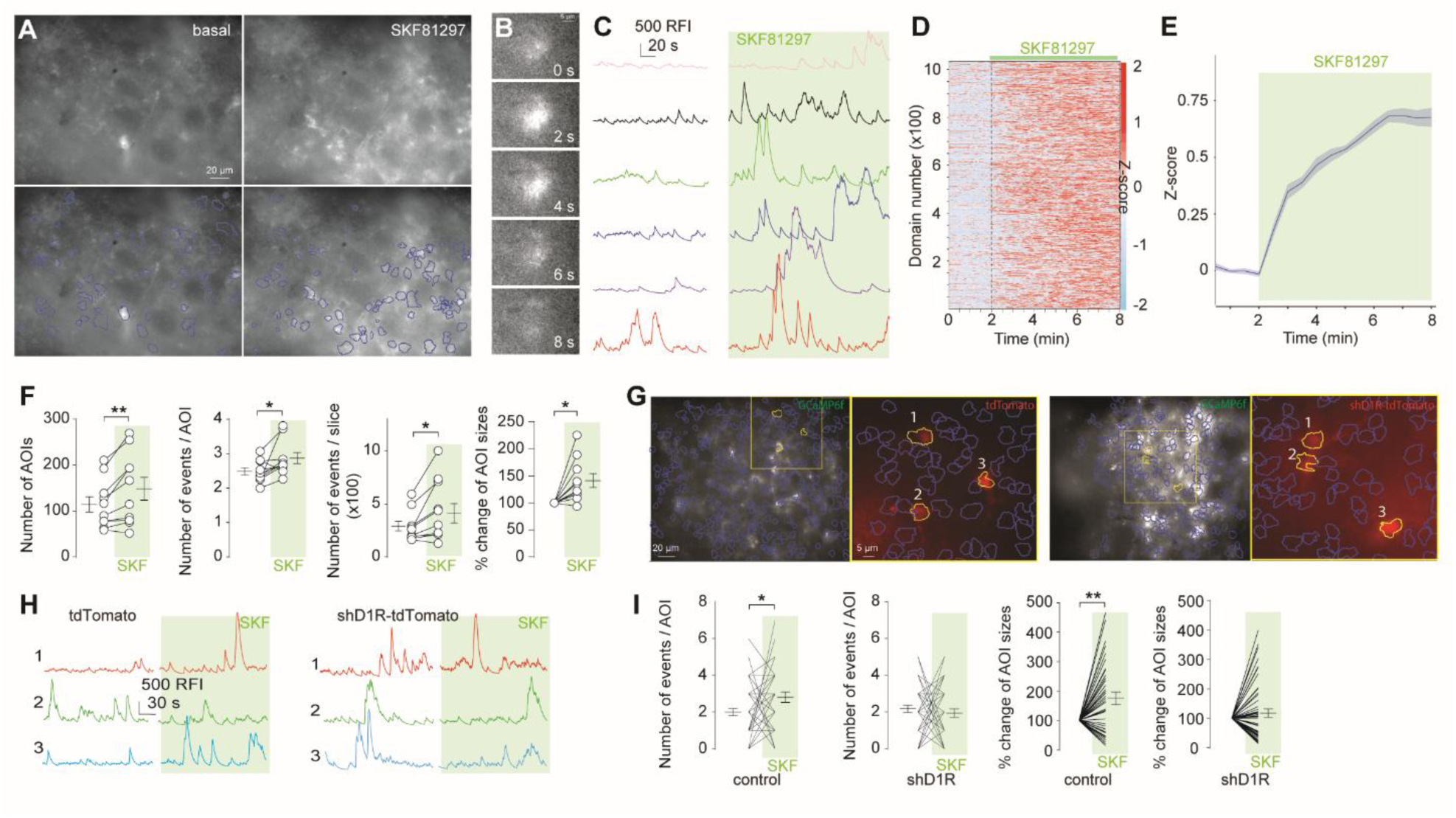
Activation of D1R increases Ca^2+^ activity in NAcSh astrocytes. (A) Example 3-min stacked images of GCaMP6f-mediated fluorescence in NAcSh slices through the GFP channel during perfusion with aCSF (upper left) or SKF81297 (upper right). Locations and areas of GCaMP6f signals extracted from the same slice shown above during perfusion with aCSF (lower left) or SKF81297 (lower right). (B) Example images of a single AOI showing the temporal dynamics of a Ca²⁺ activity event. (C) Ca²⁺ transients from example AOIs during perfusion with aCSF or SKF81297. (D) Heatmap showing the Z-scored astrocytic Ca²⁺ fluorescence signals from individual AOIs before and during SKF application. Signals were binned into 0.5-s intervals and Z-scored per AOI. (E) Time course of the mean Z-scored astrocytic Ca^2+^ fluorescence signals before and during SKF81297 application. (F) Summaries demonstrating that acute perfusion of SKF81297 increased the number of AOIs exhibiting Ca^2+^ events (before, 114.6 ± 16.8; SKF, 147.9 ± 24.7; *t*_9_ = 3.5, *p* < 0.01), the average number of Ca^2+^ events in each AOI during a 3-min imaging period (before, 2.5 ± 0.1; SKF, 2.9 ± 0.2; *t*_9_ = 2.4, *p* = 0.04), the total number of Ca^2+^ events across all AOIs during the 3-min imaging period (before, 283.7 ± 46.2; SKF, 441.8 ± 91; *t*_9_ = 3, *p* = 0.02), and the sizes of AOIs, calculated as the total area swept by all Ca²⁺ transients, normalized to the 3-min baseline average and expressed in % (SKF, 141.4 ± 12.8; *t*_9_ = 3.3 *p* = 0.01). n=10 slices from 4 mice, Paired two-tailed t tests. (G) Example images of Ca^2+^ fluorescence signals and AOIs extracted from slices with GfaABC1D AAV-mediated expression of tdTomato and shD1R-tdTomato (black-white), and magnified images showing overlaps between AOIs and cells expressing tdTomato or shD1R-tdTomato (red). (H) Ca²⁺ transients from example AOIs extracted from slices expressing tdTomato or shD1R-tdTomato. (I) Summaries showing that perfusion of SKF81297 increased the number of Ca²⁺ events (before, 2 ± 0.2; SKF, 2.8 ± 0.3; *t*_43_ = 2.2, *p* = 0.03) and the percentage change in the size (SKF, 177.9 ± 19%; *t*_42_ = 4.10, *p* < 0.01) of AOIs overlapping with tdTomato-expressing astrocytes, but these changes were not observed in AOIs overlapping with shD1R-tdTomato-expressing astrocytes (events before vs. SKF 2.2 ± 0.2 vs. 2 ± 0.2, *t*_50_ = 0.6, *p* = 0.56; size before vs. SKF 116.1 ± 14.1%, *t*_47_ = 1.1, *p* = 0.26). n = 43-51 AOIs obtained from 10 slices/4-5 mice/group. Paired two-tailed *t* tests. \**p* < 0.05; \*\**p* < 0.01.

Z-score analysis revealed that the overall intensity of fluorescence transients across all AOIs increased during SKF81297 perfusion (Fig. 3C-E). Detailed analysis of individual AOIs showed that D1R activation increased the total number of AOIs, the frequency of Ca^2+^ events within each AOI, the total event count per slice, and the mean size of individual AOIs (Fig. 3F). Thus, D1R activation stimulates intracellular Ca^2+^ activity in NAcSh astrocytes.

A critical question is whether the above astrocytic Ca^2+^ dynamics were mediated by astrocytic D1R or resulted from neuronal influence. While the use of TTX ruled out action potential-mediated neuronal input, it did not exclude action potential-independent processes. To examine a cell-autonomous role for D1R in astrocytes, we selectively expressed shD1R-tdTomato in NAcSh astrocytes, using the expression of tdTomato alone as controls. Only tdTomato+ astrocytic AOIs were selected for analysis (Fig. 3G), and Ca^2+^ activity within these AOIs was compared before versus after SKF81297 application, with pre-application activity serving as the baseline for each AOI. SKF81297 application increased both the frequency of Ca^2+^ events and the size of AOIs in control tdTomato+ astrocytes, whereas no significant changes were observed in shD1R-tdTomato+ AOIs (Fig. 3I). Taken together, these results support the hypothesis that astrocytic D1R plays a cell-autonomous role in elevating Ca^2+^ signaling within NAcSh astrocytes.

### Astrocytic D1R is required for cocaine-induced morphometric changes of NAcSh astrocytes

In rats following abstinence from cocaine self-administration, the overall surface area and volume of astrocytes in the NAc core are reduced, and these changes are accompanied by reduced expression of GFAP and reduced colocalization of astrocytic processes with synaptic markers (29). Such morphometric changes of NAc astrocytes are thought to impair glutamate homeostasis and drive maladaptive synaptic changes that increase vulnerability to cocaine taking and relapse (30–32).

Using astrocyte-specific viral expression of either tdTomato alone (control) or shD1R-tdTomato, we measured structural remodeling in the mouse NAcSh. We found that the surface area of control NAcSh astrocytes decreased following both 1-d and 45-d abstinence from cocaine self-administration (Fig. 4A, B). However, in astrocytes expressing shD1R, this cocaine-induced reduction in surface area was abolished (Fig. 4A, B).

**Figure 4.**
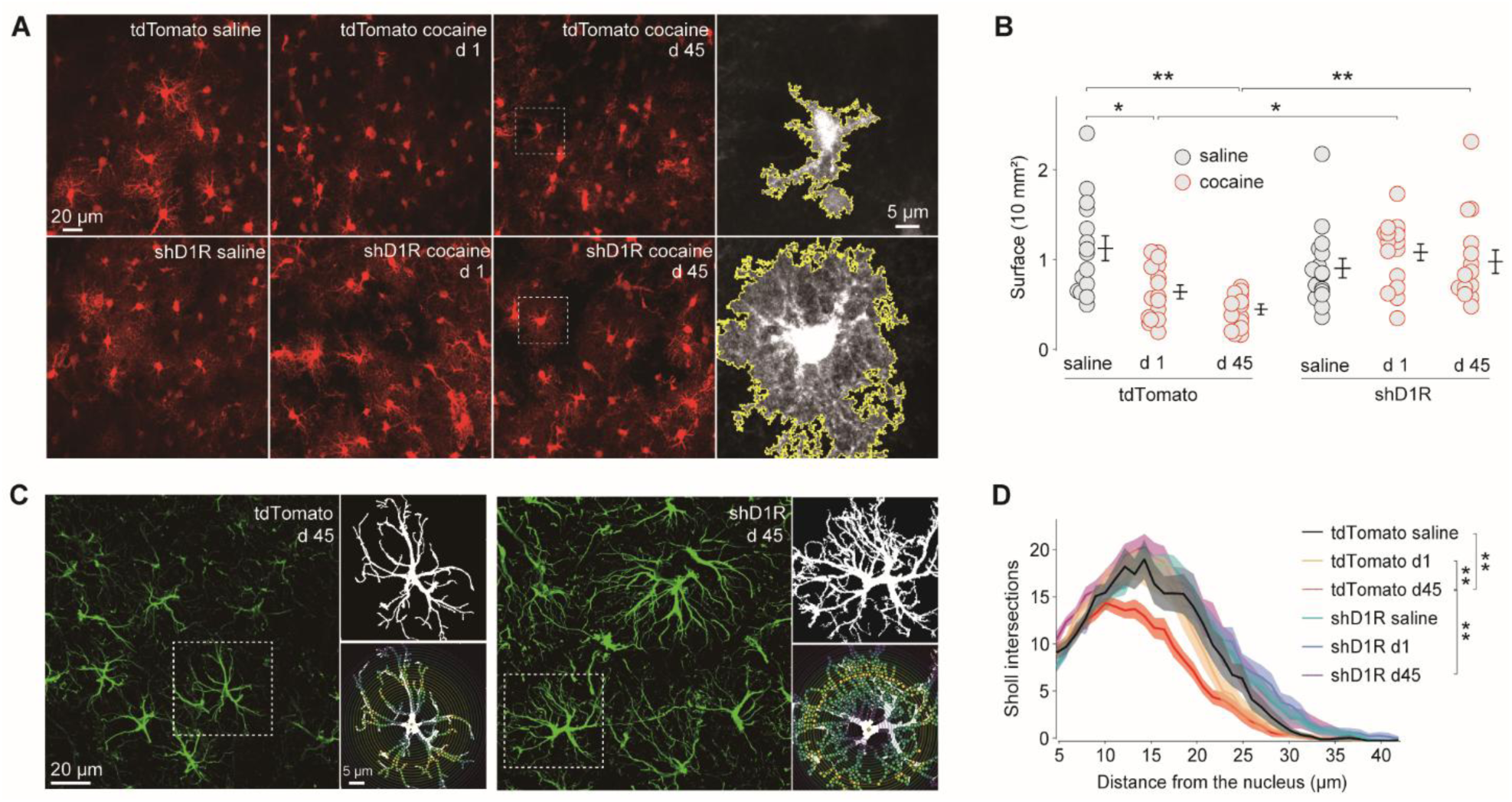
Knockdown of astrocytic D1R prevents cocaine-induced morphometric changes of NAcSh astrocytes. (A) Example images of NAcSh slices from mice injected with AAV5-GfaABC1D-tdTomato or AAV5-GfaABC1D-shD1R-tdTomato showing astrocytic morphology following saline or cocaine self-administration at abstinence day 1 and day 45. Right panels show enlarged single-astrocyte images with cell boundaries outlined (yellow) for morphological visualization. (B) Summaries (each dot representing one astrocyte) showing reduced surface areas of NAcSh astrocytes at abstinence day 1 and day 45 in tdTomato control mice but not in shD1R mice (in mm^2^: tdTomato saline, 1.1 ± 0.1, n = 15/3; tdTomato d1, 0.6 ± 0.1, n = 16/4; tdTomato d45, 0.5 ± 0.1, n = 17/4; shD1R saline, 0.9 ± 0.1, n = 16/4; shD1R d1,1.1 ± 0.1, n = 16/4; shD1R d45, 1.0 ± 0.1, n = 15/3; *F*_5,89_ = 6.9, *p* < 0.01, One-way ANOVA ; *p* = 0.02 tdTomato saline vs. tdTomato d1, *p* < 0.01 tdTomato saline vs. tdTomato d45, *p* = 0.04 tdTomato d1 vs. shD1R d1, *p* < 0.01 tdTomato d45 vs. shD1R d45, Bonferroni posttests). (C) Example GFAP immunohistostaining of NAcSh slices from tdTomato and shD1R-tdTomato mice at abstinence d45 following cocaine self-administration. Dashed boxes indicating regions shown at higher magnification. Right panels showing reconstructed single astrocytes and the corresponding Sholl analyses. (D) Plots from Sholl analysis of reconstructed astrocytes expressing tdTomato or shD1R-tdTomato from mice following saline or cocaine self-administration and abstinence. Curves representing the number of Sholl intersections as a function of distance from the soma; shades indicating SEM (tdTomato saline, 7.8 ± 1.1, n = 13/3; tdTomato d1, 7.2 ± 1.2, n = 16/3; tdTomato d45, 5.6 ± 0.9, n = 21/4; shD1R saline, 8.4 ± 1.1, n = 16/3; shD1R d1, 9.0 ± 1.1, n= 17/4; shD1R d45, 9.1 ± 1.2, n = 17/3) Two-way repeated-measures ANOVA revealed a significant main effect of group (*F*_5,3384_ = 73.6, *p* < 0.01) and a significant group × distance interaction (*F*_175,3384_ = 2.3, *p* < 0.01), followed by Bonferroni-corrected posttests. *p* < 0.01 tdTomato saline vs. tdTomato d45, *p* < 0.01 tdTomato d1 vs. tdTomato d45, *p* < 0.01 tdTomato d45 vs. shD1R d45. \**p* < 0.05, \*\**p* < 0.01.

Furthermore, the Sholl intersection analysis, used to quantify the complexity of astrocytic branching, revealed a decrease in the number of processes in control astrocytes after 45 days of cocaine abstinence. This decrease in structural complexity was also prevented by the knockdown of astrocytic D1R (Fig. 4C, D). Additional Sholl-based metrics, including the mean and maximum numbers of intersections, showed a similar reduction in control mice following cocaine abstinence, and these reductions were abolished in shD1R-expressing astrocytes (Fig. S2E, F).

Importantly, the expression of shD1R alone did not alter the surface area or process number in saline mice (Fig. 4B, D). These findings suggest that, rather than maintaining basal astrocytic morphology, astrocytic D1R-signaling is selectively implicated in the adaptive changes of NAcSh astrocytes that occur during cocaine taking or abstinence.

### Astrocytic D1R contributes to cocaine-induced generation of silent synapses

Cocaine self-administration generates glutamatergic synapses that contain NMDA receptors (NMDARs) but lack functionally stable AMPA receptors (AMPARs). These so-called AMPAR-silent synapses represent new, immature synaptic contacts that mediate the initial remodeling of NAcSh circuits and promote cocaine seeking (2, 15). Application of either cocaine (12) or D1 dopamine receptor agonists (Fig. 3) to NAcSh slices increases astrocytic Ca^2+^ activity, while dampening astrocytic Ca^2+^ activity prevents cocaine-induced generation of silent synapses, tying astrocytes to this synaptogenesis process (12, 17). Given the persistent elevation of dopamine in the NAcSh following cocaine exposure, we examined whether astrocytic D1R was required for the generation of these silent synapses.

We selectively expressed shD1R-tdTomato in NAcSh astrocytes in mice, using tdTomato expression alone as controls. Following cocaine self-administration and 1 day of abstinence, we used a minimal stimulation assay to assess the percentage of silent synapses on NAcSh MSNs adjacent to tdTomato+ astrocytes (see Methods; Fig. 5A-C). In saline mice, the levels of silent synapses were similar between MSNs either adjacent to tdTomato-control or shD1R-expressing astrocytes, indicating that astrocytic D1R knockdown did not alter basal synaptic connectivity (Fig. 5D-H).

**Figure 5.**
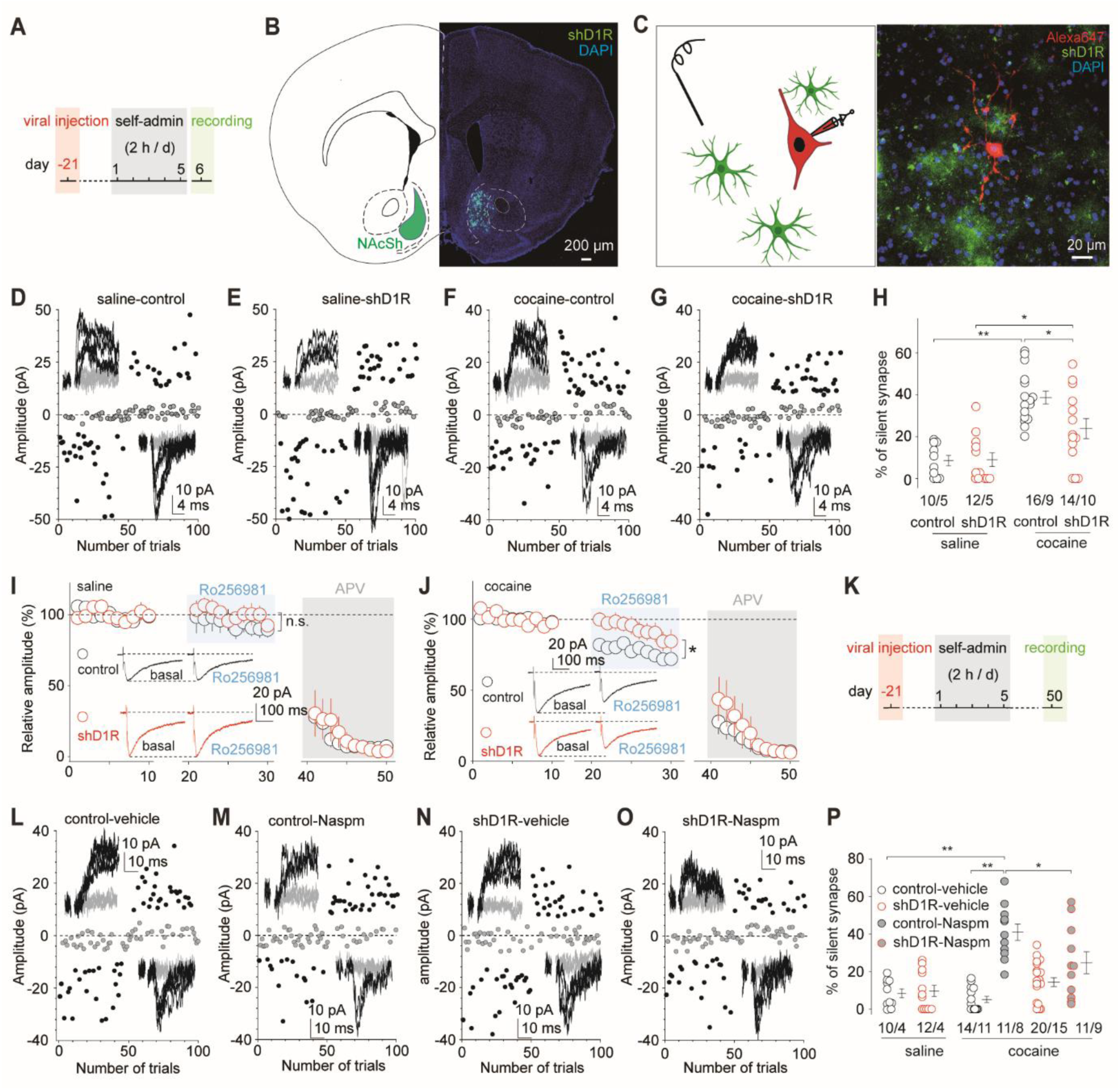
Knockdown of astrocytic D1R dampens cocaine-induced generation of silent synapses. (A) Timeline for experiments shown in (B-J). (B) Diagram and example image showing astrocyte-selective expression of shD1R-GFP in the NAcSh. (C) Diagram and example image showing an MSN (filled with red Alexa Fluor 647) adjacent to GFP-expressing astrocyte processes that was chosen for recording. (D-G) Example EPSCs evoked in NAcSh MSNs at −70 mV or +50 mV (insets) over 100 trials recorded 1 day after saline or cocaine self-administration from mice with astrocytic expression of GFP or shD1R-GFP. (H) Summaries showing that the percentage of silent synapses in MSNs was increased on d1 after cocaine in GFP mice, but the increase was attenuated in shD1R mice (in %: saline-GFP, 8. 6 ± 2.6%, n = 10/5; saline-shD1R, 9.0 ± 3.29, n = 12/5; cocaine-GFP, 38.6 ± 3.1, n = 16/9; cocaine-shD1R, 23.9 ± 4.8, n = 14/10; *F*_3,48_ = 15.8, *p* < 0.01, one-way ANOVA; *p* < 0.01 saline-GFP vs. cocaine GFP, *p* = 0.04 saline-shD1R vs. cocaine-shD1R, *p* = 0.02 cocaine-GFP vs. cocaine-shD1R, Bonferroni posttests). (I) Summaries showing that, following Ro256981 application, NMDAR EPSC amplitudes were comparable between GFP and shD1R-GFP mice 1 day after saline self-administration (relative amplitude in %: GFP, 94.1 ± 1.2, n = 9/5; shD1R, 100.5 ± 1.4, n = 8/6, main effect of group *F*_1,15_ = 0.6, *p* = 0.46, two-way ANOVA repeated measures). Subsequent application of APV (50 μM) abolished synaptic currents, confirming that they were mediated by NMDARs. Inset: example NMDAR EPSCs before and during Ro256981 application. (J) Summaries showing that Ro256981 inhibited the amplitude of NMDAR EPSCs after cocaine self-administration, with a significantly smaller reduction in shD1R-GFP mice compared to GFP controls (relative amplitude in %: GFP, 77.7 ± 1.3, n = 9/4; shD1R, 92.7 ± 2.0, n = 9/4, main effect of group *F*_1,16_ = 5.6, *p* = 0.03, two-way ANOVA repeated measures). (K) Timeline for experiments shown in (L-P). (L-O) Example EPSCs recorded in the absence or presence of Naspm at −70 mV or +50 mV over 100 trials in MSNs from GFP or shD1R mice 45 days after cocaine self-administration. (P) Summaries showing that while Naspm increased the percentage of silent synapses in both GFP and shD1R-GFP mice, this Naspm-mediated increase was attenuated in shD1R mice (in %: saline-GFP, 8.4 ± 2.3, n = 10/4; saline-shD1R, 9. 7 ± 3.0, n = 12/4; cocaine-GFP-vehicle, 5.1 ± 1.7, n = 14/11; cocaine-GFP-Naspm, 41.1 ± 4.4, n = 11/8; cocaine-shD1R-vehicle, 14.4 ± 2.4, n = 20/15; cocaine-shD1R-Naspm, 24. 8 ± 5.8, n = 11/9, *F*_5,72_ = 14.8, *p* < 0.01, one-way ANOVA; *p* < 0.01 saline-GFP vs. cocaine-GFP-Naspm, *p* < 0.01 cocaine-GFP-vehicle vs. cocaine-GFP-Naspm, *p* = 0.03 cocaine-GFP-Naspm vs. cocaine-shD1R-Naspm, Bonferroni posttests). \**p* < 0.05, \*\**p* < 0.01.

Consistent with previous findings (11–15), we observed that cocaine self-administration increased the proportion of silent synapses in control NAcSh MSNs (Fig. 5H). However, this cocaine-induced generation of silent synapses was dampened in MSNs adjacent to shD1R-expressing astrocytes (Fig. 5D-H). Notably, D1R knockdown did not completely abolish the effect; a modest increase in silent synapses remained detectable in cocaine-trained shD1R mice (Fig. 5H). This partial effect may be attributable to incomplete knockdown or suggest that astrocytic D1R is only one component of a broader signaling network mediating cocaine-induced synaptogenesis.

Cocaine-induced synaptogenesis involves a coordinated sequence of events, including the insertion of GluN2B-containing NMDA receptors into new synaptic sites (3, 8, 17, 33–35). To determine whether astrocytic D1R influences cocaine-induced synaptogenesis upstream of this step, we recorded NMDAR-mediated excitatory postsynaptic currents (EPSCs) in MSNs. We isolated NMDAR EPSCs at a holding potential of -40 mV (to relieve Mg^2+^-mediated blockade) in the presence of the AMPAR-selective antagonist NBQX (5 μM). In saline mice, NMDAR EPSCs in both control and shD1R preparations were resistant to exposure to the GluN2B-NMDAR-selective antagonist Ro256981 (200 nM), indicating low basal synaptic expression of GluN2B NMDARs and that this low expression was not affected by astrocytic D1R knockdown (Fig. 5I). In cocaine-trained mice, a portion of NMDAR EPSCs was inhibited in MSNs adjacent to control astrocytes (Fig. 5J), confirming that silent synapse generation is accompanied by the synaptic recruitment of GluN2B NMDARs. However, this recruitment was attenuated in MSNs adjacent to shD1R-expressing astrocytes (Fig. 5J), suggesting that the astrocytic D1R-signaling acts upstream of GluN2B NMDAR insertion.

Following their generation, a portion of silent synapses mature by recruiting Ca^2+^-permeable AMPARs (CP-AMPARs), rendering them no longer ‘silent’. These unsilenced synapses persist during prolonged drug abstinence to stabilize the remodeled NAcSh circuit (15). These cocaine-generated synapses can be unmasked by the pharmacological inhibition of CP-AMPARs (11). We predicted that, since shD1R impairs the initial generation of silent synapses, there would be a corresponding reduction in matured, cocaine-specific synapses after long-term abstinence. After 45 days of abstinence from cocaine administration, we measured silent synapse levels in the presence of the CP-AMPAR-selective antagonist Naspm (200 μM). In control MSNs (without astrocytic shD1R), levels of silent synapses were low but elevated upon perfusion of Naspm, revealing the unsilenced cocaine-generated synapses (Fig. 5L-P). In MSNs adjacent to shD1R-expressing astrocytes, however, the level of Naspm-unmasked synapses was lower than in controls (Fig. 5L-P), suggesting a reduced number of cocaine-generated synapses.

Taken together, these results demonstrate that astrocytic D1R is a key component of the astrocyte-neuron signaling network that initiates cocaine-induced synaptogenesis. By impairing the initial generation of silent synapses, astrocytic D1R knockdown may compromise long-term circuit remodeling associated with cocaine memories.

### Astrocytic D1R knockdown reduces cocaine seeking

Formation of new synapses is a fundamental mechanism for redefining neural circuit connectivity that underlies new behaviors. Silent synapse-mediated remodeling of NAcSh circuits has been implicated in the development of pathophysiological motivational states induced by drugs of abuse (2, 8, 15, 16). Having established a critical role of astrocytic D1R in silent synapse generation and subsequent maturation, we next examined whether this astrocytic signaling pathway can be experimentally targeted to reduce drug seeking.

We selectively expressed shD1R-tdTomato or tdTomato alone as a control in NAcSh astrocytes. Following 5 days of cocaine self-administration, mice began 10 days of 2-h daily extinction training starting on abstinence day 2. This was followed by 3 days of tests for cue-induced reinstatement of cocaine seeking (Fig. 6A). shD1R and control mice exhibited similar levels of lever-pressing during the cocaine self-administration period, indicating that astrocytic D1R knockdown did not impair the primary operant learning or the reinforcing effects of cocaine (Fig. 6B).

**Figure 6.**
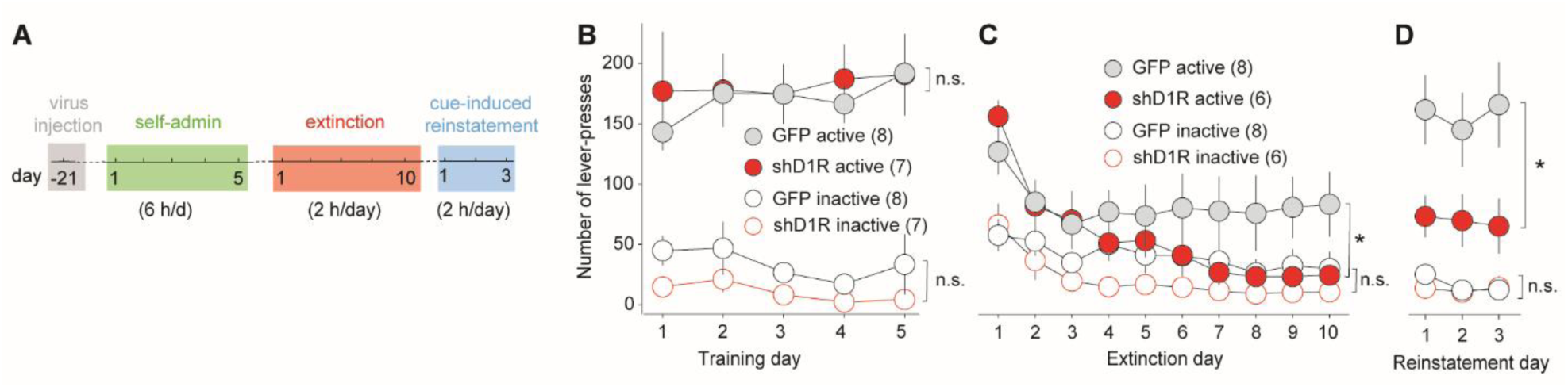
Knockdown of astrocytic D1R expedites extinction of cocaine seeking and decreases cue-induced reinstatement of cocaine seeking. (A) Schematic showing experimental procedure. (B) Summaries showing that GFP and shD1R-GFP mice exhibited comparable cocaine self-administration acquisition (active lever-press: time × group interaction *F*_2.2, 29_ = 0.4, *p* = 0.68; inactive lever-press: time × group interaction *F*_1.9, 24.7_ = 0.2, *p* = 0.84, two-way ANOVA repeated measures). (C) Summary of lever-pressing responses on the last day of the 10-day extinction procedure (day 10) in GFP and shD1R-GFP mice (active lever-press on d10: GFP, 86.1 ± 26, n = 8 vs. shD1R-GFP, 21.5 ± 5.6, n = 6, *t*_7.6_ = 2.4, *p* = 0.04; inactive lever-press: GFP, 27.9 ± 11.4, n = 8 vs. shD1R-GFP, 11.2 ± 7.8, n = 6, *t*_11.5_ = 1.2, *p* = 0.25, two-tailed Welch’s *t* tests). (D) Summaries showing lever-press responses during cue-induced reinstatement tests across three days (active lever-press: main effect of group *F*_1,12_ = 5.7, *p* = 0.04; inactive lever-press: main effect of group *F*_1,12_ = 0.3, *p* = 0.6, two-way ANOVA repeated measures). \**p*<0.05.

During extinction training, active lever-presses no longer resulted in cocaine delivery or the presentation of cocaine-associated cues. tdTomato control mice showed resistance to extinction, maintaining high levels of operant responding over the 10-d period despite the absence of reinforcement (Fig. 6C). In contrast, shD1R mice exhibited accelerated extinction kinetics, with a rapid decline in the operant response within the first few days of training (Fig. 6C). Furthermore, during the subsequent cue-induced reinstatement tests, shD1R mice showed a marked reduction in lever-pressing compared to controls.

Thus, compromising astrocytic D1R-signaling, a manipulation that impairs silent synapse generation and the subsequent remodeling of NAcSh circuits, undermined the persistence of the cocaine- and cue-associative memories that drive drug seeking.

## Discussion

While it has long been known that dopamine receptors are expressed in astrocytes, their cellular and behavioral roles remain underexplored. Our current study demonstrates that knockdown of astrocytic D1R dampens cocaine-induced generation of silent synapses, thereby compromising the subsequent remodeling of NAcSh circuits. Consequently, the selective knockdown of D1R in NAcSh astrocytes reduces resistance to cocaine extinction and attenuates cue-induced reinstatement of cocaine seeking. These results reveal astrocytic D1R as a key cellular substrate through which cocaine experience induces long-lasting circuit and behavior alterations related to SUD.

### Astrocytic D1R is a key component of NAcSh dopamine signaling

Dopaminergic signaling in the NAcSh regulates several forms of synaptic plasticity that are critical for the development of SUD-related behaviors (1, 2). While many of these effects are attributable to neuronal D1R, there are also experimental observations that cannot be explained by neuronal D1R alone. For example, global knockout of D1R impairs several forms of synaptic plasticity, but it remains a puzzle why inhibition of D1R influences NAcSh neurons that do not express D1R (36). Our current results show that astrocytic D1R is a key component of NAcSh dopaminergic signaling during cocaine-induced synaptogenesis. These results provide a novel angle to understanding the role of dopaminergic signaling in circuit remodeling and behavioral alterations related to SUD.

The NAcSh is one of a few brain regions where astrocytes are dopaminoceptive, expressing relatively high levels of dopamine receptors (37–39). We found that the probability of astrocytic Ca^2+^ events increases following perfusion of D1R agonists, an effect attenuated by D1R knockdown. These findings align with previous reports of increased astrocytic Ca^2+^ activity following optogenetic stimulation of dopaminergic presynaptic terminals or cocaine application (12, 22). Importantly, while D1R is often regarded as Gs-coupled, D1R-mediated astrocytic Ca^2+^ dynamics require IP3R2, suggesting a Gq-signaling pathway (22, 40, 41).

Recent results have begun to link astrocytic D1R to NAcSh physiology, such as the rapid release of adenosine and acute inhibition of presynaptic activity (22). Behaviorally, mice with astrocytic D1R knockout exhibit reduced locomotor response to acute administration of amphetamine (22). These results, taken together with our current findings, establish astrocytic D1R-signaling as an important component underlying the most fundamental functions of dopamine: its instructive roles in neural plasticity and learning (42–44).

### Astrocytic D1R in cocaine-induced synaptogenesis

Cocaine-induced generation of silent synapses is a complex synaptogenic process, involving concerted neuronal and nonneuronal mechanisms (15, 17). Exposure to cocaine increases the activity of transcription factor CREB in both NAc MSNs and astrocytes (35, 45). Upregulation of CREB activity in MSNs is mediated, in part, by neuronal D1R-Gs-signaling pathway, which is essential for cocaine-induced synapto- and spinogenesis (2, 4, 16, 45). Upregulation of astrocytic CREB activity leads to increased glutamatergic synaptic transmission to MSNs (35), with its role in synaptogenesis remaining to be determined.

In parallel, cocaine self-administration mobilizes an astrocyte-mediated developmental mechanism: astrocytes release thrombospondins, which, in turn, interact with neuronal _α_2_δ_-1, considered an auxiliary subunit of voltage-gated Ca^2+^ channels, to promote the formation of new synapses (12, 17). However, it remains unclear how these astrocytic responses are induced by cocaine experience. Our current findings indicate that astrocytic D1R is a potential mediator. Specifically, cocaine-induced dopamine accumulation and the resulting repeated activation of D1R-Gq signaling increase the Ca^2+^ dynamics in NAcSh astrocytes (Fig. 3) (12). It has been demonstrated in the somatosensory cortex that increased astrocytic Ca^2+^ dynamics lead to thrombospondin release (46). Conversely, preventing Ca^2+^ dynamics in NAcSh astrocytes prevents cocaine-induced synaptogenesis (12). Thus, the D1R-Gq-Ca^2+^ pathway in NAcSh astrocytes may serve as a signaling route through which cocaine experience triggers astrocytic release of thrombospondins to promote synaptogenesis in this brain region.

Our results show that astrocytic knockdown of D1R compromised, but did not abolish, cocaine-induced generation of silent synapses (Fig. 5). This partial effect may be associated with the efficacy of astrocytic D1R knockdown. Specifically, although shD1R decreased D1R mRNA by ∼60% (Fig. S1), residual astrocytic D1R could still support a low level of the D1R-coupled synaptogenic signaling in response to cocaine. Furthermore, although we selected MSNs adjacent to shD1R-tdTomato signals, there might be other, uninfected astrocytes nearby that provide D1R-coupled synaptogenic signaling. Moreover, the astrocytic D1R-signaling pathway likely co-exists with other pathways, such as astrocytic glutamate- or serotonin-signaling, which are simultaneously and cooperatively operating to promote synapse formation. In this case, disrupting one pathway can only reduce but not eliminate the synaptogenic effect of cocaine.

### Astrocytic D1R in cocaine seeking

During cocaine self-administration training, two parallel learning processes operate simultaneously: (1) instrumental learning, through which operant responding is acquired and driven primarily by the reinforcing effect of cocaine; and (2) associative learning, through which conditioned stimuli (CS) acquire reinforcing properties that promote operant responding even in the absence of cocaine.

While both learning processes involve dopaminergic signaling in the NAcSh, excitotoxic lesions of this region has yielded inconsistent and often limited effects on the acquisition of drug self-administration (47–51), suggesting that NAcSh plays a sub-essential role in instrumental learning. Consequentially, it is unsurprising that cocaine-reinforced operant responding remained unaltered following the manipulation of NAcSh astrocytic D1R (Fig. 6).

In contrast, during extinction and reinstatement tests (Fig. 6), operant responses are not reinforced by cocaine. Instead, they are driven by CS, such as contextual, discrete or internal cues, that become reinforcing through associative learning during cocaine self-administration training. It has been hypothesized that cocaine-generated synapses encode CS-cocaine associative memories that support CS-induced drug seeking (2, 15). Supporting this hypothesis, disrupting the generation or maturation of silent synapses in the NAcSh decreases cue-induced cocaine seeking without affecting the initial acquisition of cocaine self-administration (11–14). Thus, astrocytic D1R knockdown may compromise CS-cocaine associative memories by influencing silent synapse generation or maturation, resulting in the observed decrease in operant responding in both extinction and reinstatement.

Unlike rats, mice well trained with cocaine self-administration exhibit operant responses that are highly resistant to instrumental extinction (52–54). In our study, the number of lever-presses in control mice remained persistently high over 10 days of training (Fig. 6C). This persistence may stem from the high endurance of drug-seeking memories, the formation of which may be undermined by impaired circuit connectivity remodeling following astrocytic D1R knockdown. Alternatively, this persistence may reflect inefficient extinction learning in mice. Extinction is not simple erasure of existing memories but also a process involving the formation of new inhibitory memories (55, 56). The restoration of astrocytic morphologies and neuron-astrocyte contacts following astrocytic D1R knockdown (Fig. 4) may recover the astrocyte’s capacity to support extinction-related synaptic remodeling, improving extinction learning. In shD1R mice, operant responses during the first three days of extinction were similar to controls but declined in later sessions (Fig. 6C). This pattern suggests the possibility that, while basal instrumental memories remain intact, the efficiency of extinction learning is enhanced in shD1R mice.

In the reinstatement test, contingent light cues elicited high levels of operant responding in control mice, indicating strong cue-cocaine associative memories. The decreased responding in shD1R mice may reflect compromised associative memories. However, because shD1R mice entered the reinstatement test following more effective instrumental extinction, the reduction in responses is likely multi-factorial, involving both weakened original memories and enhanced formation of extinction memories.

Activation of D1R increased Ca^2+^ dynamics in NAcSh astrocytes, potentially activating Ca^2+^-dependent signaling pathways that initiate astrocyte-neuron communication (Fig. 3). Possibly due to a relatively low basal dopamine tone, D1R knockdown did not alter the basal properties of astrocytes and their adjacent MSNs (Figs. 2, 5). However, during the dramatic increase in dopamine tone induced by cocaine, astrocytic D1R is strongly and repeatedly activated. Under these conditions, D1R knockdown prevented morphometric changes in astrocytes and dampened the generation of silent synapses (Figs. 4, 5). These cellular changes correlate with expedited cocaine extinction and reduced cue-induced reinstatement (Fig. 6). These results position the astrocytic D1R as a detector of dopamine dynamics that regulate neural circuit remodeling, offering a novel cellular target for reducing drug-induced behaviors.

## Methods

Our study examined male and female animals, and similar findings are reported for both sexes. Primary astrocyte cultures were also prepared from mice of both sexes.

### Animals

Wild-type C57BL/6J mice were purchased from The Jackson Laboratory (Bar Harbor, ME) and used for behavioral, electrophysiological, morphological, and molecular experiments. Aldh1l1-Cre/ERT2 × ROSA26-Lck-GCaMP6f-flox mice were used for astrocytic Ca²⁺ imaging studies. Mice were 6–8 weeks old at the start of experimental procedures and were housed under a 12-h light/dark cycle with food and water available ad libitum.

### Viral vectors

For astrocytic D1R knockdown, recombinant adeno-associated viral vectors (AAVs) were used to express a microRNA-adapted short hairpin RNA (shRNAmir) targeting the mouse dopamine D1 receptor gene (Drd1). The shRNA target sequences (bwD1a: 5′-ACCGATGTCTCTCTAGAAA-3′) and (bwD1b: 5’-CCACTTTGTATTTATGTAA-3’) were embedded in two concatenating miR-30 backbones with a bulge in the stem, cloned into the 3’UTR of the viral vector plasmid (12), and expressed under the control of the astrocyte-selective GfaABC1D promoter. AAVs included AAV-GfaABC1D-shD1R-GFP, AAV-GfaABC1D-shD1R-tdTomato, AAV-GfaABC1D-GFP, and AAV-GfaABC1D-tdTomato (titer ≥ 4.1 × 10¹² viral genomes/ml). All viruses were packaged as serotype 5 by the University of North Carolina Gene Therapy Center Vector Core.

### Primary astrocyte cultures

Primary astrocyte cultures were prepared from the striatum of postnatal day 0 (P0) or P1 mice. Dissection and dissociation were performed as previously described (57). A total of 10^6^ cells per well in a poly-D-lysine-coated 6-well plate were plated into 5% Serum Medium, MEM with Earle’s salts (Millipore Sigma M4655), supplemented with 5% fetal bovine serum and 0.2% MITO+ Serum-Extender (Corning, #355006). Cells were transduced at 7 days in vitro (DIV) with 4 µl AAV and harvested on DIV28.

### Semiquantitative RT-PCR

Primary astrocytes transduced with AAVs were washed with PBS with Ca^2+^ and Mg^2+^ and lysed in Trizol (Thermo-Fisher). RNA was isolated following the manufacture’s protocol. Using random primers, cDNA synthesis was performed as described previously(57). PCRs were performed for mouse Drd1 with the intron flanking primers (5’-GAGTCCAGGGGTTTTGGGAG-3’) and (5’-GGCTTAGCCCTCACGTTCTT-3’), and for GAPDH with the intron flanking primers (5’-TCAGGAGAGTGTTTCCTCGTC -3’) and (5’-GATGGGCTTCCCGTTGATGA-3’).

### Single-nucleus RNA sequencing

Bilateral NAcSh tissues were collected from cocaine- or saline-trained mice (n = 6 per group), snap-frozen on dry ice, and stored at −80 °C until use. Single-nucleus isolation was performed following the 10x Genomics protocol (CG000124, Isolation of Nuclei for Single Cell RNA Sequencing), as previously described (58). snRNA-seq libraries were prepared using the GEM-X Universal 3′ Gene Expression v4 kit (10x Genomics). Isolated nuclei and barcoded gel beads, together with reverse transcription reagents, were loaded into the Chromium X system to generate single-nucleus gel bead-in-emulsions (GEMs). Following cDNA amplification and cleanup, libraries were constructed through cDNA fragmentation, end repair and A-tailing, adaptor ligation, and sample index PCR. Library concentrations were quantified using a Qubit 4.0 fluorometer (Life Technologies), and library quality was assessed using a TapeStation 2200 (Agilent Technologies). Pooled libraries were sequenced on an Illumina NextSeq 2000 using paired-end sequencing (Read 1: 28 bp; i7 index: 10 bp; i5 index: 10 bp; Read 2: 90 bp), with an average sequencing depth of ∼30,000 reads per nucleus.

### Single-nucleus RNA-seq data analysis

Raw sequencing data were processed using Cell Ranger (v9.0.0; 10x Genomics) for sample demultiplexing, barcode processing, UMI counting, and alignment to the mouse reference transcriptome (GRCm39-2024-A). The resulting UMI count matrices were imported into Seurat (v5.3.0) (59) in R (v4.5.1) for downstream analysis. Nuclei were excluded if they contained fewer than 300 detected genes, fell within the top 2% of total UMI counts, or contained more than 15% mitochondrial gene content. Doublets were identified and removed using scDblFinder (v1.22.0) (60). After quality control and filtering, 11,029 nuclei from saline-treated mice and 17,611 nuclei from cocaine-treated mice were retained for analysis. Gene expression data were log-normalized using the Seurat function NormalizeData function and scaled using ScaleData function. The top 2,000 highly variable genes were identified using FindVariableFeatures function with the “vst” method and used to compute the first 30 principal components. Clustering was performed using the Seurat function FindClusters function. Differential gene expression between conditions was assessed using FindMarkers function with the MAST test (test.use = “MAST”), requiring a minimum expression in 20% of nuclei (min.pct = 0.2), an adjusted p value < 0.05, and a log fold-change threshold of 0.38.

### In vivo virus injection

Mice were anesthetized by intraperitoneal injection of a ketamine (100 mg/kg)–xylazine (10 mg/kg) mixture and placed in a stereotaxic frame (Stoelting Co., Wood Dale, IL). Using a 10-μL NanoFil syringe fitted with a 32-gauge needle and controlled by a UMP3 Micro4 pump system (World Precision Instruments, Sarasota, FL), AAV solutions were bilaterally infused into the NAcSh at a rate of 200 nL/min (500 nL per hemisphere; coordinates relative to bregma: AP +1.60 mm, ML ±0.70 mm, DV −4.45 mm). The needle was left in place for an additional 5 min after infusion to allow diffusion and minimize backflow.

### Catheter implantation

Catheter implantation for cocaine self-administration was performed as previously described (61–63). Briefly, a silastic catheter was advanced approximately 1 cm into the right jugular vein, and the distal end was tunneled subcutaneously to exit at the back between the scapulae. Catheters were constructed from silastic tubing (length, 7 cm; inner diameter, 0.30 mm; outer diameter, 0.64 mm) and connected to a Vascular Access Button (Instech). Mice were allowed to recover for at least 3 days following surgery, during which general health and body weight were monitored. During the recovery period, catheters were flushed daily with sterile saline containing cefazolin (3 mg/mL) and heparin (3 U/mL) to prevent infection and maintain catheter patency.

### Self-administration apparatus

All behavioral experiments were conducted in operant conditioning chambers (Med Associates). Each chamber (15.24 × 13.34 × 12.7 cm) was equipped with one active and one inactive lever, a conditioned stimulus (CS) light positioned above the active lever, and a house light. No food or water was available in the chamber during training or testing sessions.

### Intravenous cocaine self-administration

Cocaine self-administration training began at least 3 days after surgery. During the 5-day training period, mice were placed in the operant chamber for daily 6-hour sessions. Presses on the active lever resulted in an infusion of cocaine (1 mg/kg delivered over 3-6 s) and simultaneous illumination of the CS light above the lever and the house light. Both the CS light and house light remained illuminated for 20 s, during which additional lever presses were recorded but did not result in further cocaine infusions. Presses on the inactive lever had no reinforced consequences but were recorded. After the 5-day training period, mice were returned to their home cages and remained there during the abstinence period. Cocaine-trained mice that failed to meet the self-administration criteria (≥60 infusions per session and ≥70% of total lever presses on the active lever) were excluded from further experimentation and analysis. For electrophysiological recordings on abstinence day 1, mice were used 1 day after the final training session. For recordings on abstinence day 45, mice were singly housed in their home cages throughout the 44-day abstinence period, with food and water available ad libitum.

### Cocaine extinction and cue-induced reinstatement of cocaine seeking

The extinction procedure consisted of 2-h daily sessions conducted for 10 consecutive days. During each session, mice were placed in the same operant chambers used for cocaine self-administration. Active lever presses did not result in cocaine infusion or presentation of CS cues. Inactive lever presses were recorded but had no programmed consequences. Cue-induced reinstatement tests consisted of 2-h daily sessions conducted for 3 consecutive days. During each session, mice were placed in the same operant chambers. Active lever presses resulted in presentation of the light cue previously paired with cocaine delivery, but without cocaine infusion. Inactive lever presses had no programmed consequences.

### Preparation of acute brain slices

Mice were decapitated under deep isoflurane anesthesia. Coronal brain slices (250 μm thick) containing the nucleus accumbens (NAc) were prepared using a VT1200S vibratome (Leica) in ice-cold cutting solution containing (in mM): 135 N-methyl-D-glucamine, 1 KCl, 1.2 KH₂PO₄, 0.5 CaCl₂, 1.5 MgCl₂, 20 choline bicarbonate, and 11 glucose. The solution was saturated with 95% O₂ / 5% CO₂, adjusted to pH 7.4 with HCl, and osmolality was adjusted to 305 mmol kg⁻¹. Slices were then transferred to artificial cerebrospinal fluid (aCSF) containing (in mM): 119 NaCl, 2.5 KCl, 2.5 CaCl₂, 1.3 MgCl₂, 1 NaH₂PO₄, 26.2 NaHCO₃, and 11 glucose, with osmolality adjusted to 285–295 mmol kg⁻¹. Slices were incubated in oxygenated aCSF (95% O₂ / 5% CO₂) at 37°C for 30 min and subsequently maintained at 20–22°C for at least 30 min before experimentation.

### Electrophysiological recordings

Whole-cell patch-clamp recordings were performed to examine membrane properties of astrocytes and synaptic transmission onto NAcSh MSNs. All recordings were obtained from the ventromedial NAcSh.

For synaptic transmission recordings, MSNs closely adjacent to astrocytic GFP fluorescence were preferentially selected. During recordings, slices were superfused with aCSF preheated to 30–32°C using a feedback-controlled in-line heater (Warner). To measure minimal stimulation–evoked responses and NMDAR-mediated EPSCs, recording electrodes (2–5 MΩ) were filled with a Cs⁺-based internal solution containing (in mM): 135 CsMeSO₃, 5 CsCl, 5 TEA-Cl, 0.4 EGTA (Cs), 20 HEPES, 2.5 Mg-ATP, 0.25 Na-GTP, and 1 QX-314 (Br), pH adjusted to 7.3. Picrotoxin (0.1 mM) was included in the aCSF during all recordings to inhibit GABA_A_ receptor-mediated currents. Presynaptic afferents were stimulated using a constant-current isolated stimulator (Digitimer) through a monopolar electrode (glass pipette filled with aCSF). Series resistance was typically 7–20 MΩ, left uncompensated but monitored continuously. Cells exhibiting changes in series resistance >20% were excluded from analysis. Synaptic currents were recorded using a MultiClamp 700B amplifier, filtered at 2.6–3 kHz, amplified fivefold, and digitized at 20 kHz.

To record NMDAR-mediated EPSCs, the AMPAR-selective antagonist NBQX (5 μM) and picrotoxin (0.1 mM) were included in the bath solution. The membrane potential was held at −40 mV to relieve Mg²⁺-dependent block of NMDARs. After adjusting stimulation position and intensity, recording conditions remained constant throughout the experiment. Following a 10-min baseline recording, a bath solution containing the GluN2B-selective antagonist Ro256981 (200 nM) was perfused into the recording chamber. Given the wash-in time and use-dependent nature of Ro256981, EPSCs recorded 10 min after antagonist application were used to assess Ro256981-sensitive NMDAR currents. After 20-min recording in Ro256981 bath, a bath containing APV (50 μM) instead of Ro256981 was applied to confirm that the recorded EPSCs were NMDAR-mediated.

Silent synapses were assessed using the minimal stimulation assay as previously described (3, 11, 34, 64). Briefly, after obtaining small (∼50 pA) EPSCs at −70 mV, stimulation intensity was gradually reduced until synaptic failures and successes could be clearly distinguished. Stimulation intensity and frequency were then held constant for the remainder of the experiment. For each neuron, 50–100 traces were recorded at −70 mV and +50 mV, and recordings were repeated for one to two additional rounds. Only neurons exhibiting stable failure rates (changes <15% between rounds) were included in the analysis. Failures and successes were visually scored in a blinded manner to minimize bias.

The percentage of silent synapses was calculated based on two assumptions: (1) presynaptic release sites are independent, and (2) release probability is identical across all synapses, including silent synapses. Under these assumptions, silent synapse percentage was calculated as:

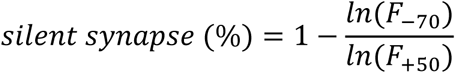

where F_−70_ and F_+50_ represent failure rates at −70 mV and +50 mV, respectively (64). Even when these assumptions were not strictly met, this equation was still used to qualitatively predict changes in silent synapse levels, as previously validated (13).

To assess electrophysiological properties of astrocytes, whole-cell recordings were obtained from NAcSh astrocytes identified by GFP expression driven by rAAV5-GfaABC1D-GFP. Recording electrodes (4–6 MΩ) were filled with a K⁺-based internal solution containing (in mM): 135 K-gluconate, 5 KCl, 0.4 EGTA, 10 HEPES, 2.5 Mg-ATP, and 0.3 Na-GTP (pH 7.3, adjusted with KOH), supplemented with 0.1% Dextran–Texas Red. Cells were held at −80 mV, and voltage steps from −120 mV to +40 mV were applied in 10-mV increments for 500 ms. Steady-state currents were measured during the final portion of each voltage step.

### Imaging astrocytic Ca^2+^ activity in slices

To image Ca²⁺ activity in NAcSh astrocytes, GCaMP6f was expressed in astrocytes by crossing Aldh1l1-Cre/ERT2 transgenic mice with ROSA26-Lck-GCaMP6f-flox-GFP reporter mice (The Jackson Laboratory, stock nos. 029655 and 029626). Offsprings of both sexes, aged 8–16 weeks, received five daily intraperitoneal injections of tamoxifen (100 mg/kg, dissolved in corn oil; VWR, #9001-30-7) to induce astrocyte-specific GCaMP6f expression.

Three weeks after the last tamoxifen injection, coronal brain slices (200-μm thick) containing NAcSh were prepared, initially incubated at ∼34°C for 30 min, and subsequently kept at room temperature (∼22°C) in aCSF, saturated with 95% O2 and 5% CO2. All recordings were performed in the continuous presence of TTX (1 μM). GCaMP6f fluorescence signals were captured using a Hamamatsu ORCA-ER C4742 CCI camera mounted on an Olympus BX51W1 microscope through the 488-nm filter. The camera was set with a fixed exposure time of 500 ms and a frame speed of 2 fps. The temperature of the imaging chamber was set at ∼31°C. The imaging was confined to the medial ventral portion of the NAcSh through a 40× water immersion objective lens. The sampling area of the slice was 36,000 μm^2^ (1344 × 1024 pixels). To generate heatmaps and time courses of astrocytic Ca²⁺ activity (z-score), baseline GCaMP6f fluorescence was recorded for 2 min, followed by perfusion with SKF81297 (1 μM) for an additional 6 min. In some experiments, GCaMP6f fluorescence was measured repeatedly, with each recording lasting 3 min and separated by a 3-min interval.

For experiments involving astrocytic shD1R expression and the corresponding controls, double-transgenic mice (Aldh1l1-Cre/ERT2 x ROSA26-Lck-GCaMP6f-flox-GFP) received intra-NAcSh injections of either AAV-GfaABC1D-shD1R-tdTomato or AAV-GfaABC1D-tdTomato. One week after viral injection, intraperitoneal tamoxifen administration was initiated, and Ca²⁺ imaging experiments were performed three weeks after the completion of tamoxifen treatment. To select the areas of interest (AOI) originating from astrocytes infected by either shD1R AAV or control AAV, extracted AOIs were overlaid with the tdTomato fluorescence images, and those overlapping with tdTomato signals were included for further analysis.

Image analysis was performed using both ImageJ and Python. Fluorescence signals were pre-processed in ImageJ using the rolling ball algorithm to remove uneven background illumination across frames. Each video was then subjected to motion correction, source extraction, and deconvolution in Python using the Calcium Imaging Analysis (CaImAn) software package. Motion correction was performed using the NoRMCorre algorithm, and fluorescence signals were further processed with a high-pass filter. A modified version of the constrained nonnegative matrix factorization framework (CNMF-E), specialized for de-mixing and de-noising one-photon imaging data, was used to extract fluorescent signals. The spatial-temporal features (including shapes, locations, and peaks of Ca²⁺ transients) extracted by CNMF-E were subsequently used to identify and classify fluorescent events.

To isolate astrocytic activities and minimize noise and neuropil contamination, only fluorescent events with a signal-to-noise ratio (SNR) greater than 2 were retained for further analysis. A customized Python script was developed to analyze Ca²⁺ transients. Ca²⁺ transients were included in the analysis if their peak amplitudes were at least twofold higher than the baseline standard deviation and separated by at least 1000 ms. Ca²⁺ transients were plotted based on the signals extracted by CNMF-E. The number of Ca²⁺-mediated event locations was determined by counting the number of AOIs, where fluorescent events exceeded an SNR threshold of 2. Each AOI was considered to represent a separate astrocyte domain. The number of events per AOI was calculated by averaging the number of Ca²⁺ transients detected in each AOI within the field of view. The number of events per slice was determined by counting the total number of Ca²⁺ transients across the entire recording period. The percentage change in AOI sizes was calculated as the total net changes in fluorescence signals over the baseline, multiplied by the duration of the transients, and then normalized to the mean baseline fluorescence intensity and expressed as a percentage.

### Immunohistochemistry and confocal microscopy

Mice were transcardially perfused with 0.1 M sodium phosphate buffer, followed by 4% paraformaldehyde (PFA) in 0.1 M phosphate buffer. Brains were removed and postfixed in 4% PFA overnight at 4°C. Whole brains were then stored in 30% sucrose in phosphate buffer with 0.1% sodium azide at 4°C until sectioning. Coronal sections (30-μm thick) containing the NAc were cut on a microtome (Leica) and collected in 4°C PBS. For neuronal labeling, sections were incubated with mouse anti-NeuN antibody (1:500; Cell Signaling Technology, #94403). For astrocyte labeling, sections were incubated with rabbit anti-GFAP antibody (1:500; Agilent/Dako, #Z0334). Following primary antibody incubation, sections were washed three times in PBS (5 min each) and incubated for 2 h at room temperature with appropriate fluorophore-conjugated secondary antibodies (1:300). Alexa Fluor 594–conjugated donkey anti-mouse IgG (Invitrogen, Thermo Fisher Scientific, #A-21203) was used for NeuN staining. For GFAP staining, Alexa Fluor 488–conjugated or Alexa Fluor 594–conjugated donkey anti-rabbit IgG (Invitrogen, Thermo Fisher Scientific, #A21206 or #A21207) was used as indicated for different experiments. Sections were washed again 3 × 5 min in PBS before they were mounted in ProLong Gold Antifade with DAPI (Molecular Probes).

Sections were imaged using a Leica TCS SP5 confocal microscope controlled by Leica Application Suite software. Images were acquired with a 5× air objective and 40× or 60× oil-immersion objectives. Whole-slice images were obtained using automated tile scanning followed by image stitching.

### Astrocyte morphological analysis

Astrocyte surface area was quantified from confocal images using Fiji (ImageJ, NIH). Z-stack images of tdTomato-labeled astrocytes in the NAcSh were acquired using identical imaging settings across all experimental groups. Maximum intensity projections were generated for analysis. Individual astrocytes with clearly identifiable somas and non-overlapping processes were manually selected. Cell boundaries were traced using the freehand selection tool based on tdTomato fluorescence, and the enclosed area was measured using the “Measure” function in Fiji. All measurements were performed by an experimenter without the knowledge of mice’ treatments.

Sholl analysis was performed to quantify the morphological complexity of astrocytes. Individual astrocytes were reconstructed from confocal z-stack images using ImageJ/Fiji. Images were first background-subtracted and thresholded, and individual astrocytes were manually isolated to exclude overlapping processes from neighboring cells. Binary images were then generated and skeletonized prior to analysis. Sholl analysis was conducted using the Sholl Analysis plugin in ImageJ/Fiji. Concentric circles were drawn at 1-μm intervals centered on the soma of each astrocyte, and the number of process intersections at each radial distance from the soma was quantified. Sholl intersection profiles were averaged across cells within each experimental group and plotted as a function of distance from the soma.

### Drugs and reagents

Cocaine hydrochloride (HCl), provided by the NIDA Drug Supply Program, was dissolved in 0.9% NaCl saline. Ketamine and xylazine, purchased from a Drug Enforcement Administration–designated vendor at the University of Pittsburgh, were mixed for anesthesia. All other chemicals were purchased from Sigma-Aldrich, Tocris, J.T. Baker, Fluka, or Alomone Labs.

### Statistics

All results are presented as mean ± SEM. Data collection was randomized. Sample sizes were determined based on previous studies using similar experimental approaches (65, 66). All data were analyzed offline, with investigators blinded to experimental conditions during analysis.

A total of 305 mice were used in this study, of which 95 were excluded from the final data analysis and interpretation for the following reasons: (1) 37 mice were excluded due to post-surgical health issues (for example, a >20% reduction in body weight); (2) 41 mice were excluded due to catheter failure or failure to meet self-administration criteria; and (3) 17 mice were excluded due to off-target stereotaxic injections or poor virus expression. Repeated experiments within the same experimental group were pooled for statistical analysis.

For electrophysiological experiments, sample sizes are presented as n/m, where n represents the number of cells and m represents the number of mice. For behavioral experiments, sample sizes are presented as n, representing the number of mice. Statistical significance was assessed using unpaired two-tailed t-tests or one-way or two-way ANOVA, as specified in the relevant figure legends or texts. Statistical significance was set at p < 0.05 for all analyses. Statistical analyses were performed using GraphPad Prism (v.8).

### Study approval

All animal experiments were performed in accordance with protocols approved by the Institutional Animal Care and Use Committee at the University of Pittsburgh.

## Data availability

Values for all data points in graphs are reported in the Supporting Data Values file.

## Author contributions

SY, QL, YW, WC, ZC, AKZ, PH, CW, EJN, OMS, and YD designed research studies. SY, QL, YW, WC, ZC, AKZ, PH, ZL, XQ, ZX, YH, and GS conducted experiments and acquired data. SY, QL, YW, WC, ZC, AKZ, PH, XQ, CW, and OMS analyzed data. CW, EJN, OMS, and YD supervised the study. SY, QL, and YD wrote the manuscript. SY, QL, YW, WC, ZC, AKZ, PH, ZL, XQ, ZX, YH, GS, CW, EJN, OMS, and YD reviewed and edited the manuscript. Co–first authorship order was determined on the basis of overall contribution to the study.

## Acknowledgments

We thank Min Li for technical support. Cocaine was provided by the NIH NIDA Drug Supply Program.

