## Supplementary Materials for "Astrocytic D1 Dopamine-Signaling Regulates Synaptic Remodeling and Cocaine Seeking"

Shi Yan et al.

Figs. S1

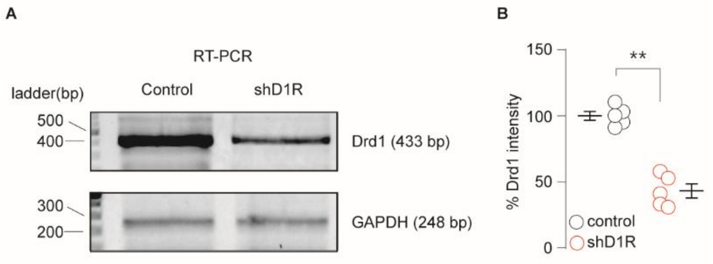


**Fig. S1. Validation of Drd1 knockdown by RT-PCR in cultured cells.**

(A) Representative RT-PCR gel images showing Drd1 expression in control and shD1R groups. GAPDH was used as an internal control. (B) Quantification of Drd1 band intensity expressed as a percentage of the control group (set as 100%). Data are presented as mean ± SEM (Control,100 ± 3.4, n= 6; shD1R, 43.2 ± 5.3, n=5, $t_{6.7} = 9$, $p<0.01$, unpaired Welch's *t* test). Each data point represents an independent culture. Statistical significance was assessed using an unpaired two-tailed *t* test. $^{**}p < 0.01$.

Figs. S2

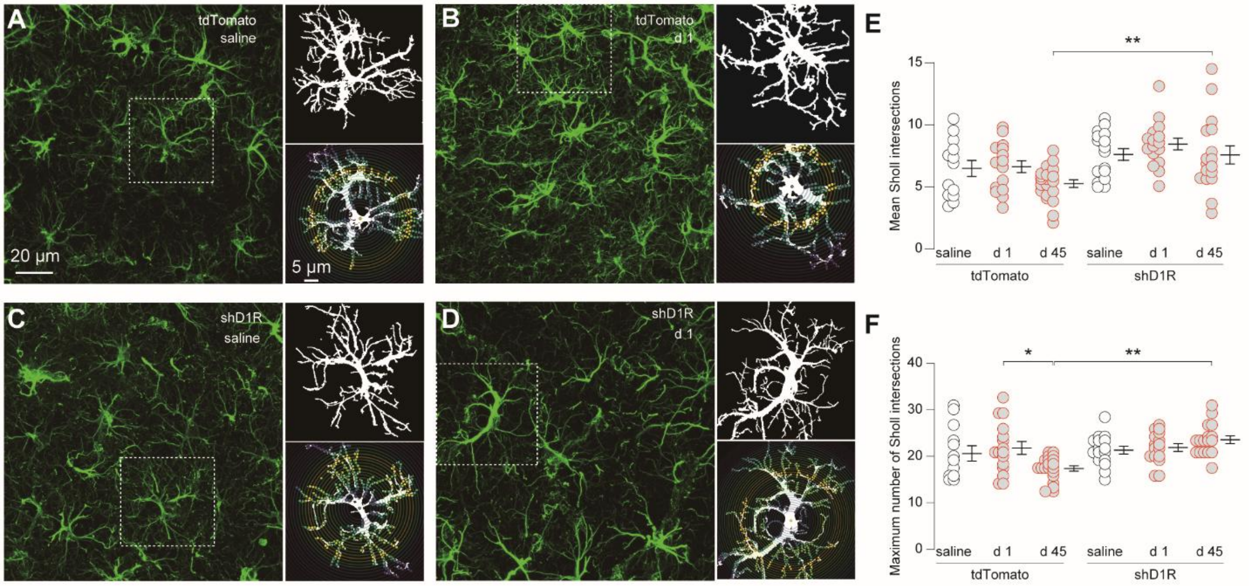


**Figs. S2 Additional Sholl analysis of NAcSh astrocytes across experimental conditions.**

(A–D) Representative GFAP immunohistostaining of NAcSh slices from tdTomato and shD1R-tdTomato mice at abstinence day1 following saline or cocaine self-administration. Dashed boxes indicate regions shown at higher magnification. Right panels show reconstructed single astrocytes and the corresponding Sholl analyses. (E) Quantification of mean Sholl intersections, calculated as the average number of intersections across all concentric radii for each reconstructed astrocyte at abstinence day 1 and day 45 (tdTomato-saline, 6.5 ± 0.7, n = 13/3; tdTomato -d1, 6.6 ± 0.5, n = 16/3; tdTomato -d45, 5.3 ± 0.3, n = 20/4; shD1R-saline, 7.6 ± 0.5, n = 16/3; shD1R-d1, 8.5 ± 0.5, n = 16/4; shD1R-d45, 7.6 ± 0.7, n = 17/3. $F_{5,92} = 5.0$, $p < 0.01$, one-way ANOVA; $p = 0.02$ tdTomato -d45 vs. shD1R-d45, Bonferroni posttests). (F) Quantification of maximum Sholl intersections, defined as the peak number of intersections observed across all radii for each astrocyte at abstinence day 1 and day 45 (tdTomato-saline, 20.7 ± 1.9, n = 13/3; ctrl-d1, 22.1 ± 1.6, n = 16/3; tdTomato -d45, 16.9 ± 0.7, n = 20/4; shD1R-saline, 21.6 ± 1, n = 16/3; shD1R-d1, 22.3 ± 1, n = 16/4; shD1R-d45, 24.2 ± 1, n = 17/3. $F_{5,92} = 5.0$, $p < 0.01$, one-way ANOVA; $p = 0.02$ tdTomato-d1 vs. tdTomato-d45; $p < 0.01$ tdTomato-d45 vs. shD1R-d45, Bonferroni posttests). Each dot represents one astrocyte. Data are presented as mean ± SEM. *$p < 0.05$, **$p < 0.01$.
